# Novel diversity of polar Cyanobacteria revealed by genome-resolved metagenomics

**DOI:** 10.1101/2023.02.03.526606

**Authors:** Igor S. Pessi, Rafael Vicentini Popin, Benoit Durieu, Yannick Lara, Valentina Savaglia, Beatriz Roncero-Ramos, Jenni Hultman, Elie Verleyen, Wim Vyverman, Annick Wilmotte

## Abstract

Benthic microbial mats dominated by Cyanobacteria are important features of polar lakes. Although culture-independent studies have provided important insights into their diversity, only a handful of genomes of polar Cyanobacteria have been sequenced to date. Here, we applied a genome-resolved metagenomics approach to data obtained from Arctic, sub-Antarctic, and Antarctic microbial mats. We recovered 22 unique metagenome-assembled genomes (MAGs) of Cyanobacteria, most of which are only distantly related to genomes that have been sequenced so far. These include i) lineages that are common in polar microbial mats such as the filamentous taxa *Pseudanabaena, Leptolyngbya, Microcoleus/Tychonema*, and *Phormidium*; ii) the less common taxa *Crinalium* and *Chamaesiphon*; iii) an enigmatic Chroococcales lineage only distantly related to *Microcystis*; and iv) an early branching lineage in the order Gloeobacterales that is almost exclusively restricted to the cold biosphere, for which we propose the name *Candidatus* Sivonenia alaskensis. Our results show that genome-resolved metagenomics is a powerful tool for expanding our understanding of the diversity of Cyanobacteria, especially in understudied remote and extreme environments.

**Data summary:** The sequencing data generated in this study have been submitted to the European Nucleotide Archive (ENA) under the BioProject PRJEB59431. Individual accession numbers for raw reads and genomic bins are listed in **Table S1** and **Table S3**, respectively. Genomic bins can also be downloaded from doi.org/10.6084/m9.figshare.22003967. The commands used throughout this study are available in github.com/igorspp/polar-cyanobacteria-MAGs.

**Impact statement:** Cyanobacteria are photosynthetic microorganisms that play important roles in polar lacustrine ecosystems. Many Cyanobacteria are difficult to grow in the laboratory, particularly in isolation from other organisms, which makes it challenging to sequence their genomes. As such, considerably fewer genomes of Cyanobacteria have been sequenced so far compared to other bacteria. In this study, we used a metagenomics approach to recover novel genomes of Cyanobacteria from Arctic and Antarctic microbial mats without the need to isolate the organisms. The community DNA was extracted and sequenced, and the genomes of individual populations were separated using bioinformatics tools. We recovered 22 different genomes of Cyanobacteria, many of which have not been sequenced before. We describe in more detail an interesting lineage of ancestral Cyanobacteria in the order Gloeobacterales, for which we propose the name *Candidatus* Sivonenia alaskensis. Our study shows that genome-resolved metagenomics is a valuable approach for obtaining novel genomes of Cyanobacteria, which are needed to improve our understanding of life in the polar regions and the planet at large.

## Introduction

Microbial mats are highly successful and productive ecosystems found in a wide range of environments since the dawn of life on Earth [1, 2]. Microbial mats commonly comprise a vast diversity of microorganisms such as auto- and heterotrophic bacteria, fungi, microalgae, and heterotrophic protists embedded in an exopolysaccharide matrix [3]. Benthic microbial mats represent an important survival strategy against the harsh environmental conditions in polar and alpine lakes, and have Cyanobacteria as their primary source of organic carbon and nitrogen [4, 5]. In addition to aquatic microbial mats, Cyanobacteria are also important members of terrestrial and epi- and supraglacial communities in polar environments [6, 7].

Despite their importance, knowledge on the diversity and ecology of Cyanobacteria in polar environments is fragmentary [8]. Studies on the diversity of polar Cyanobacteria have mostly focused on microscopic identification and strain isolation [9–15], analysis of environmental 16S rRNA gene sequences [16–21], or a combination of these methods [22–24]. On one hand, the microscopic identification of Cyanobacteria is hindered by the high plasticity of taxonomic markers such as cell dimensions and division patterns and the relative paucity of morphological characters [25]. In addition, morphology-based assessments underestimate the diversity of Cyanobacteria in the environment compared to molecular approaches based on environmental DNA [19]. Molecular approaches, in turn, are hampered by the scarcity of cyanobacterial genomes stored in public databases, which are largely underrepresented compared to other microbial phyla and heavily biased towards the *Prochlorococcus/Synechococcus* clade [26, 27].

The genomic catalogue of polar Cyanobacteria is currently limited to a handful of strains, including *Pseudanabaena* sp. BC1403 and *Phormidesmis priestleyi* BC1401 from Greenland [28], *Leptolyngbya* sp. Cla-17 from the Canadian High Arctic [29], and the Antarctic strains *P. priestleyi* ULC007 [30], *Leptolyngbya* sp. BC1307 [31], *Synechococcus* sp. SynAce01 [32], and *Nostoc* sp. SO-36 [33]. Twelve other low-quality genomes obtained by a metagenome-like assembling approach of non-axenic strains are also available [34]. Genome-resolved metagenomics has been established in recent years as a powerful approach to obtain microbial genomes, as it circumvents the difficulties associated with culturing microorganisms by reconstructing microbial genomes directly from environmental DNA [35–38]. Several genomes of uncultured polar Cyanobacteria have been obtained recently using this approach, including several novel lineages of early branching Cyanobacteria in the order Gloeobacterales [39–41].

In this study, we aimed to expand the genomic catalogue of polar Cyanobacteria. To achieve this, we applied a genome-resolved metagenomics approach to data obtained from microbial mats from Arctic, sub-Antarctic, and Antarctic lakes spanning a wide geographic and limnological range. Our results include the recovery of novel genomes of polar Cyanobacteria and the description of an early branching lineage that is distributed across polar and alpine environments.

## Methods

### Sample description

We analysed 17 microbial mat samples obtained from 15 Arctic, sub-Antarctic, and Antarctic lakes **(Fig. 1, Table S1)**. The Arctic lakes are located in Svalbard and Greenland. The sub-Antarctic samples come from lakes in Macquarie Island (South Pacific Ocean) and Marion Island (South Indian Ocean), and sampling in Antarctica covered several locations in the Antarctic Peninsula, Transantarctic Mountains, and East Antarctica. The Antarctic lakes are distributed across five Antarctic Conservation Biological Regions (ACBRs) [42]. We analysed one microbial mat sample taken from the shallow region of each lake (*ca*. 0.2 m depth). In lakes Lundström and Forlidas, we analysed one additional sample taken from a deeper saline and hypersaline layer, respectively.

**Fig. 1.**
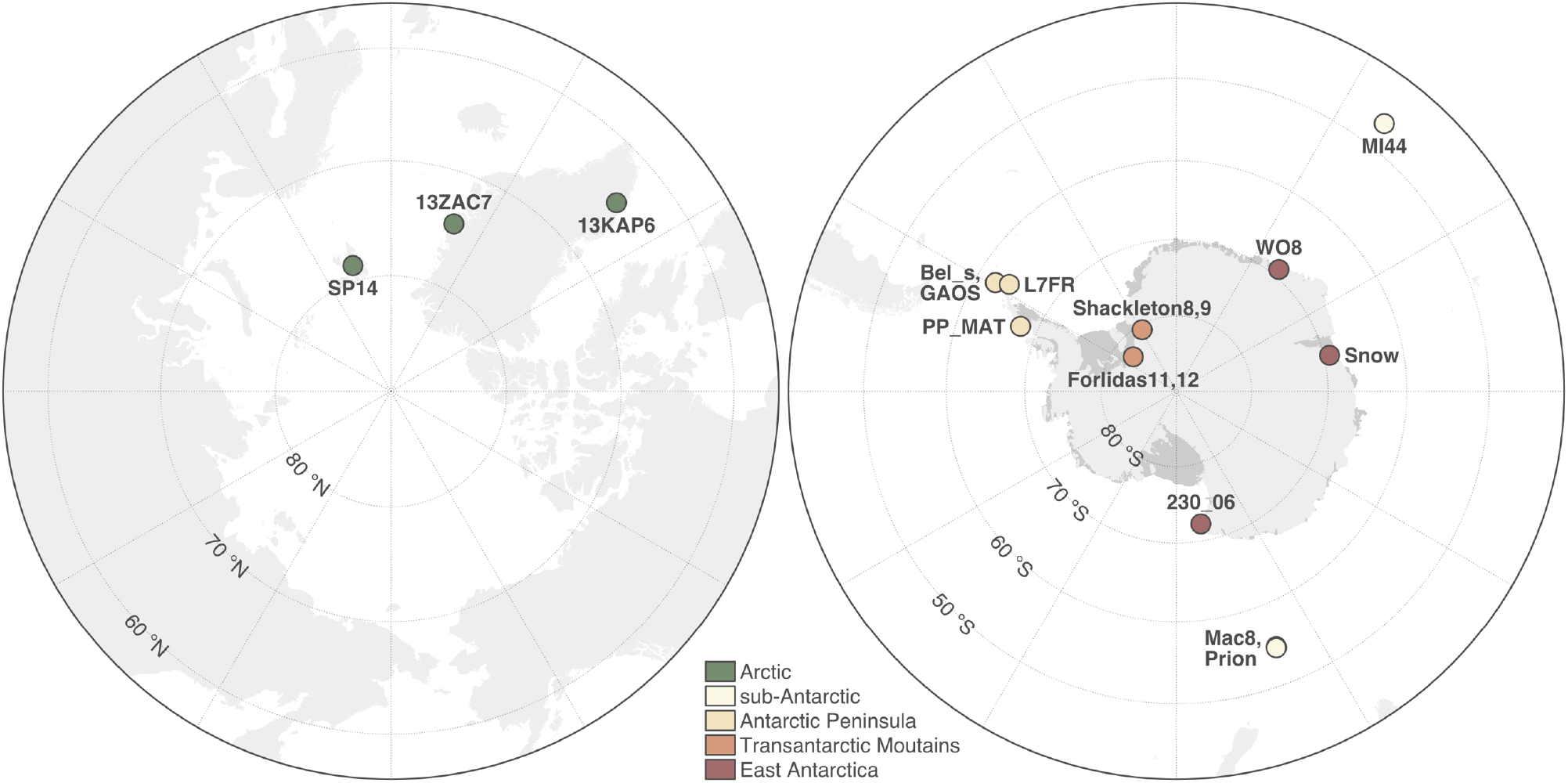
Location of the Arctic and Antarctic lakes where microbial mats were sampled. Maps were created with public data from the Norwegian Polar Institute (Tromsø, Norway). More information about the samples can be found in **Table S1**.

### DNA extraction and metagenome sequencing

We used the DNeasy PowerBiofilm DNA Isolation kit (QIAGEN, Hilden, Germany) to extract DNA from *ca*. 0.5 g of each microbial mat sample and checked the concentration and quality of the DNA extracts using the Qubit dsDNA BR Assay kit (Thermo Fisher Scientific, Waltham, MA, USA). We used the Nextera XT kit (Illumina, San Diego, CA, USA) to prepare the metagenomic libraries, which were then sent to Eurofins Genomics (Ebersberg, Germany) for sequencing using the Illumina HiSeq 2500 platform (2×100 bp). We checked the quality of the raw sequencing data with fastQC v0.11.9 (bioinformatics.babraham.ac.uk/projects/fastqc) and multiQC v1.8 [43], and used Cutadapt v1.16 [44] to trim adapters, low-quality base calls (Phred score <20), and discard short reads (<50 bp). Finally, we used METAXA v2.2 [45] to extract reads matching the 16S rRNA gene, which were then classified with mothur v1.44.3 [46] using the SILVA database release 138.1 [47] and the Naïve Bayesian Classifier with a confidence cutoff of 80% [48].

### Metagenome assembling

We assembled and binned each metagenome individually and as two co-assemblies. One co-assembly was done by grouping samples from the Arctic (n=3) and sub-Antarctic (n=3). The second co-assembly comprised the remaining samples from the Antarctic Peninsula, Transantarctic Mountains, and East Antarctica (n=11). We assembled the metagenomes with MEGAHIT v1.1.1.2 [49] and obtained 176,097 and 72,514 contigs ≥1000 bp for the Antarctic and Arctic/sub-Antarctic co-assemblies, respectively. The total assembled length was 447.6 and 182.2 Mb, respectively. The output of the individual assemblies ranged from 218 contigs/0.3 Mb (sample ‘13ZAC7’) to 96722 contigs/262.4 Mb (sample ‘PP_MAT’). The assembly of sample ‘Forlidas11’ did not yield any contig due to the very low sequencing depth achieved for this sample **(Table S1)**.

### Metagenome binning

For each individual and co-assembly, we used *anvi’o* v7.0 [50] to bin contigs ≥2500 bp into metagenome-assembled genomes (MAGs) as previously described [37, 38]. In brief, we used *Prodigal* v2.6.3 [51] to find gene calls, *HMMER* v.3.3 [52] to identify a set of 71 bacterial and 76 archaeal single-copy genes [53], and *DIAMOND* v0.9.14 [54] to assign taxonomy to the single-copy genes according to the Genome Taxonomy Database (GTDB) release 04-RS89 [55]. We used *bowtie* v2.4.2 [56] to map the quality-filtered reads from all samples to the contigs and *SAMtools* v1.1 [57] to sort and index the mapping output. We then used the *anvi-interactive* interface of *anvi’o* to manually sort the contigs into genomic bins based on differential coverage and tetranucleotide frequency. Bins that were ≥50% complete according to the presence of 71 single-copy genes [53] were manually curated using the *anvi-refine* interface of *anvi’o*. We refined the bins by removing outlying contigs according to coverage, tetranucleotide frequency, and taxonomic signal. We assigned taxonomy to the refined bins based on 122 archaeal and 120 bacterial single-copy genes with *GTDB-Tk* v1.3.0 [58] and the GTDB release 05-RS95 [55]. Bins assigned to the phylum Cyanobacteria that were ≥50% complete and ≤10% redundant – hereafter referred as MAGs – were kept for downstream analyses.

### Phylogenetic analysis

We used a concatenated alignment of 38 ribosomal proteins to place the MAGs in a phylogenetic tree alongside all genomes assigned to the Cyanobacteria/Melainabacteria group in GenBank (NCBI:txid1798711, accessed on 17 November 2022). We used *ncbi-genome-download* v0.3.1 (github.com/kblin/ncbi-genome-download) to recover the genomes from GenBank. In *anvi’o* v7.0 [50], we retrieved the translated amino acid sequence of each ribosomal protein with *HMMER* v.3.3 [52] and aligned them with *MUSCLE* v3.8.1551 [59]. We concatenated the alignments of the 38 ribosomal proteins and built a maximum-likelihood tree with *IQ-TREE* [60] using the automatic model selection and 1000 ultrafast bootstrap approximation replicates. We also used *fastANI* v1.32 [61] to calculate the genome-wide average nucleotide identity (ANI) between MAGs and GenBank genomes. For better visualization, we computed a more compact maximum-likelihood tree including only the MAGs, their closest neighbours in GenBank, strains from the Pasteur Culture Collection of Cyanobacteria (PCC), and other selected genomes. We classified the MAGs based on their phylogenetic placement following the taxonomic system of Komárek *et al*. [62].

### Gene annotation

In *anvi’o* v7.0 [50], we annotated the gene calls identified by *Prodigal* v2.6.3 [51] against the KOfam [63] and the Pfam [64] databases with *HMMER* v.3.3 [52] and the COG [65] database with *DIAMOND* v0.9.14 [54]. We also used *tblastn* (web interface, available at blast.ncbi.nlm.nih.gov/Blast.cgi?PROGRAM=tblastn&PAGE_TYPE=BlastSearch) to search for additional genes involved in mechanisms of resistance to stress. Only hits with e-value <10^−5^ and bitscore >50 were considered, according to Pearson [66].

### Distribution analyses

We used metagenomic read recruitment to compute the relative abundance of the MAGs across the 17 microbial mat samples. Prior to this, we used *dRep* v3.2.2 [67] to dereplicate the MAGs based on a 99% ANI threshold. We then used *CoverM* v0.6.1 (github.com/wwood/CoverM) to map the quality-filtered reads to the MAGs with *minimap* v2.17 [68] and compute relative abundances based on the proportion of reads recruited by the MAGs. For this, we considered only matches with ≥95% identity and ≥75% coverage. We also used *sourmash branchwater* [69, 70] and *IMNGS* [71] to search the two Gloeobacterales MAGs against metagenomic and amplicon sequencing datasets in the Sequence Read Archive (SRA), respectively. For the first, we used the *mastiff* implementation of *sourmash branchwater* (github.com/sourmash-bio/2022-search-sra-with-mastiff). The datasets where significant matches were found (containment ≥20%) were downloaded from SRA with *fasterq-dump* v3.0.1 (github.com/ncbi/sra-tools) and mapped back to the two Gloeobacterales MAGs with *CoverM* v0.6.1 as described above. For the analysis of amplicon sequencing datasets, we used the web interface of *IMNGS* (imngs.org) and only considered datasets where significant matches (≥99% similarity) accounted for ≥0.1% of the sequences.

## Results and discussion

We obtained around 500 million paired-end metagenomic reads (99.3 Gb) from 17 Arctic, sub-Antarctic, and Antarctic microbial mat samples **(Fig. 1, Table S1)**. Taxonomic profiling based on reads matching the 16S rRNA gene revealed Cyanobacteria as the second most abundant microbial phylum after Proteobacteria (mean relative abundance of 20.8 and 24.0%, respectively) **(Table S2)**. The dominance of these two phyla is commonly observed in polar microbial mats [72–74]. After assembling the reads with *MEGAHIT* [49], we used *anvi’o* [50] to manually bin and curate MAGs. Taxonomic classification based on the GTDB release 05-RS95 [55] assigned 37 MAGs to the phylum Cyanobacteria **(Fig. 2, Table 1, Table S3)**. These include two MAGs (‘PMM_0025’ and ‘PMM_0089’) belonging to the order Obscuribacterales of the Melainabacteria, a sister lineage to the Cyanobacteria *stricto sensu* (clade Oxyphotobacteria) that lacks the photosynthetic machinery [75]. Indeed, annotation of protein-coding genes revealed that the two Obscuribacterales MAGs do not encode proteins of the Calvin cycle (Rbc), photosystems I and II (Psa and Psb), cytochrome *b_6_f* complex (Pet), and phycobilisomes (Apc, Cpc, Cpe, and Pec) **(Fig. 3)**. Interestingly, the presence of genes for the small and large subunits of the nitric oxide reductase (NorC and NorB, respectively) indicates a potential role of this lineage in the production of the greenhouse gas nitrous oxide [37].

**Fig. 2.**
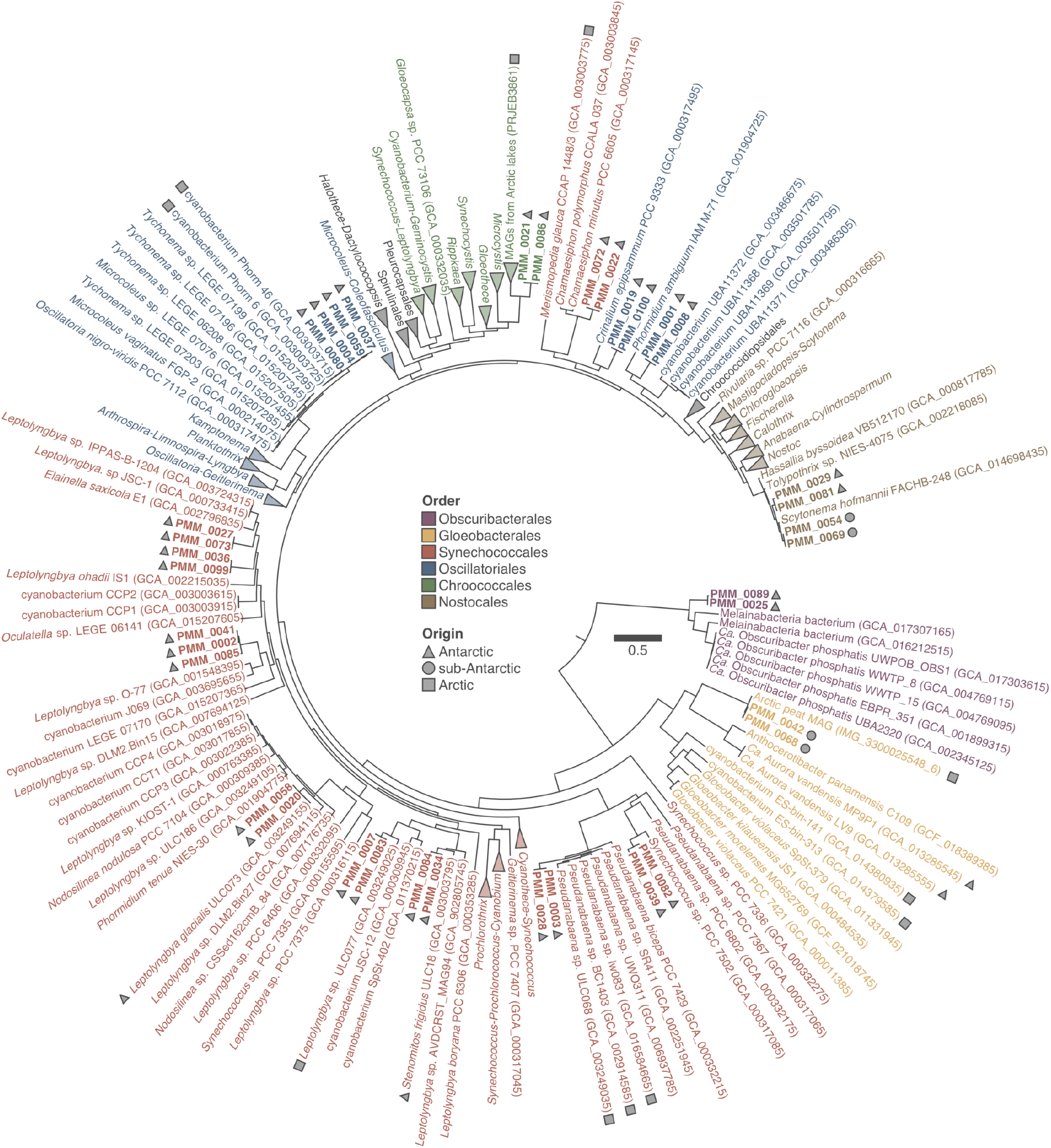
Phylogenetic analysis of 37 metagenome-assembled genomes (MAGs) assigned to the phylum Cyanobacteria, including both Cyanobacteria *stricto sensu* (clade Oxyphotobacteria) and the Melainabacteria. Maximum-likelihood tree (LG+R8 model) based on a concatenated alignment of 38 ribosomal proteins from the MAGs (in bold), their closest neighbours in GenBank, PCC strains, and other selected genomes. The geographic origin of polar MAGs and strains are indicated. Order-level classification is shown according to the taxonomic system of Komárek *et al*. [62].

**Table 1.** Information on 37 metagenome-assembled genomes (MAGs) of Cyanobacteria *stricto sensu* (clade Oxyphotobacteria) and Melainabacteria recovered from polar microbial mats.

| Group | MAG | Size (Mb) | Completion (%) | Redundancy (%) | GC (%) |
| --- | --- | --- | --- | --- | --- |
| **Melainabacteria** |  |  |  |  |  |
| Obscuribacterales | PMM_0025 | 6.1 | 90.1 | 8.5 | 47.7 |
|  | PMM_0089 | 4.7 | 77.5 | 5.6 | 47.9 |
| **Oxyphotobacteria** |  |  |  |  |  |
| Gloeobacterales | PMM_0042 | 2.9 | 97.2 | 0.0 | 49.3 |
|  | PMM_0068 | 2.8 | 95.8 | 0.0 | 48.8 |
| Synechococcales | PMM_0039 | 1.5 | 57.7 | 0.0 | 41.5 |
| (*Pseudanabaena*) | PMM_0082 | 2.7 | 85.9 | 0.0 | 40.3 |
|  | PMM_0003 | 3.1 | 78.9 | 5.6 | 42.9 |
|  | PMM_0028 | 3.7 | 84.5 | 8.5 | 42.6 |
| Synechococcales | PMM_0034 | 3.3 | 66.2 | 0.0 | 51.3 |
| (*Leptolyngbya*) | PMM_0084 | 4.8 | 81.7 | 4.2 | 51.8 |
|  | PMM_0007 | 5.2 | 94.4 | 1.4 | 49.0 |
|  | PMM_0083 | 4.6 | 94.4 | 1.4 | 48.4 |
|  | PMM_0020 | 3.2 | 90.1 | 2.8 | 57.2 |
|  | PMM_0058 | 2.8 | 84.5 | 1.4 | 56.9 |
|  | PMM_0002 | 2.9 | 90.1 | 1.4 | 52.5 |
|  | PMM_0041 | 2.5 | 50.7 | 1.4 | 52.4 |
|  | PMM_0085 | 3.7 | 81.7 | 4.2 | 52.5 |
|  | PMM_0036 | 4.6 | 66.2 | 4.2 | 49.4 |
|  | PMM_0099 | 4.7 | 67.6 | 5.6 | 49.3 |
|  | PMM_0027 | 2.2 | 73.2 | 1.4 | 55.0 |
|  | PMM_0073 | 3.6 | 85.9 | 5.6 | 55.3 |
| Synechococcales | PMM_0022 | 2.5 | 80.3 | 2.8 | 44.9 |
| (*Chamaesiphon*) | PMM_0072 | 2.9 | 83.1 | 1.4 | 44.4 |
| Oscillatoriales | PMM_0004 | 4.4 | 60.6 | 7.0 | 45.6 |
| (*Tychonema/Microcoleus*) | PMM_0037 | 5.2 | 67.6 | 8.5 | 45.3 |
|  | PMM_0059 | 4.9 | 90.1 | 5.6 | 45.6 |
|  | PMM_0080 | 6.4 | 94.4 | 2.8 | 45.1 |
| Oscillatoriales | PMM_0019 | 5.7 | 95.8 | 2.8 | 45.2 |
| (*Crinalium*) | PMM_0100 | 3.8 | 80.3 | 2.8 | 45.4 |
| Oscillatoriales | PMM_0001 | 5.8 | 94.4 | 2.8 | 45.4 |
| (*P. ambiguum*) | PMM_0008 | 5.8 | 94.4 | 2.8 | 45.4 |
| Chroococcales | PMM_0021 | 4.0 | 77.5 | 2.8 | 39.0 |
|  | PMM_0086 | 3.8 | 80.3 | 2.8 | 38.9 |
| Nostocales | PMM_0029 | 2.7 | 73.2 | 7.0 | 42.0 |
|  | PMM_0081 | 4.4 | 91.5 | 2.8 | 42.1 |
|  | PMM_0054 | 5.1 | 90.1 | 2.8 | 42.2 |
|  | PMM_0069 | 3.0 | 76.1 | 0.0 | 41.6 |

Completion and redundancy were estimated based on the presence of 71 single-copy genes [53].
More information about the MAGs can be found in **Table S3**.

**Fig. 3.**
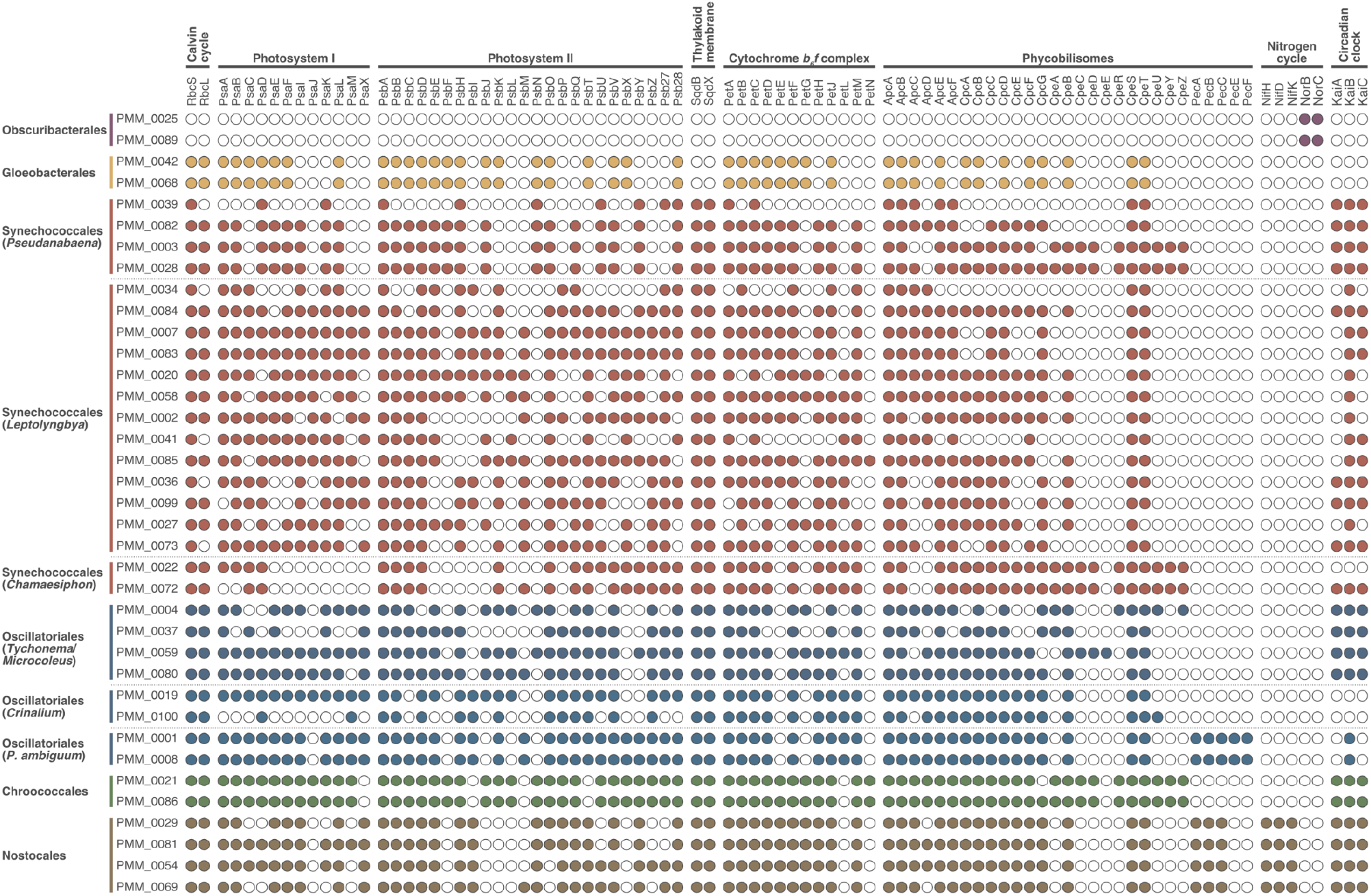
Presence of genes involved in carbon fixation, photosynthesis, nitrogen cycle, and circadian clock in 37 metagenome-assembled genomes (MAGs) of Cyanobacteria *stricto sensu* (clade Oxyphotobacteria) and Melainabacteria.

### Genome-resolved metagenomics is a reliable tool for the investigation of cyanobacterial diversity

Phylogenetic analysis based on a concatenated alignment of 38 ribosomal proteins assigned the 35 Oxyphotobacteria MAGs to five orders according to the taxonomic system of Komárek *et al*. [62]: Gloeobacterales (n=2), Synechococcales (n=19), Oscillatoriales (n=8), Chroococcales (n=2), and Nostocales (n=4) **(Fig. 2)**. Most MAGs originated from the individual (n=20) and Antarctic (n=15) (co-)assemblies **(Table S3)**. We did not recover any MAG from the individual assemblies of Arctic samples despite the high abundance of Cyanobacteria in these samples (5.9–28.6% of the reads matching the 16S rRNA gene) **(Table S2)** and the high sequencing depth (3.6–7.5 Gb) **(Table S1)**. MAG dereplication based on a 99% ANI threshold grouped the 37 Cyanobacteria MAGs into 22 unique clusters **(Table S3)**. In general, we observed a good correspondence between individual and co-assembly MAGs, *i.e*. closely related genomic bins with ≥99% ANI were recovered from the two assembly types.

The robustness of our metagenomic approach is further illustrated by the high similarity that some of the MAGs share with genomes available in GenBank. In particular, two MAGs obtained from different assemblies (‘PMM_0058’ and ‘PMM_0020’) are almost identical (99.7–99.8% ANI) to the genome of the strain *Leptolyngbya glacialis* ULC073 **(Fig. 2, Table S3)**. This is not surprising given that the three genomes originate from the same hypersaline brine layer in the benthos of Forlidas Pond, Transantarctic Mountains [24]. *L*. *glacialis* ULC073 and *L*. *antarctica* ULC047 (Ace Lake, Princess Elizabeth Land) [12], which share identical 16S rRNA gene sequences, are representative strains of an ubiquitous morphotype in Antarctic lakes belonging to the *Leptolyngbya*-*Nodosilinea* clade [12, 20, 23, 24]. Despite the importance of this lineage, the genome of *L*. *glacialis* ULC073 currently available in GenBank (accession GCA_003249155.1), which was obtained from a non-axenic unialgal culture using a metagenome-like approach [34], is very fragmented (650 contigs, N_50_=10.7 Kbp) and somewhat redundant (7.0% according to our analysis of 71 single-copy genes). Based on these parameters, the MAGs ‘PMM_0058’ and ‘PMM_0020’, which are 84.5–90.1% complete, 1.4–2.8% redundant, comprise 290–339 contigs, and have an N50 of 12.1–13.0 Kbp **(Table 1, Table S3)**, can be considered better representatives of this important lineage of Antarctic Cyanobacteria.

Other MAGs that are closely related to strains are the Nostocales MAGs ‘PMM_0054’ and ‘PMM_0069’. Genome-wide analysis revealed that they share 93.8–94.5% ANI with their closest genome on GenBank, *Scytonema hofmannii* FACHB-248 **(Fig. 2, Table S3)**. However, their 16S rRNA gene is 99.4% similar to the sequence of *Dactylothamnos antarcticus* CENA433 isolated from a freshwater biofilm in the Antarctic Peninsula [76], for which genomic information is currently lacking. The other two Nostocales MAGs (‘PMM_0029’ and ‘PMM_0081’) are also likely related to *D*. *antarcticus* given their close phylogenetic relationship with ‘PMM_0054’ and ‘PMM_0069’ **(Fig. 2)**. Finally, the Gloeobacterales MAGs ‘PMM_0042’ and ‘PMM_0068’ share 97.2% ANI with the MAG ‘IMG_3300025548_6’ recovered from peat soil in Alaska [39] **(Fig. 2, Table S3)**.

### Metagenomics reveals novel genomic diversity of polar Cyanobacteria

Phylogenetic placement and genome-wide comparison with sequences from GenBank revealed that most MAGs differ from genomes that have been sequenced so far **(Fig. 2, Table S3)**. In particular, 19 of the 37 MAGs have <80% ANI with genomes currently available in GenBank and 12 are only distantly related to existing genomes (80.1–93.2% ANI). Interestingly, phylogenetic placement clustered 16 and eight MAGs alongside polar and alpine strains, respectively **(Fig. 2)**. This is in agreement with previous studies showing that many lineages of Cyanobacteria are distributed across the cold biosphere [20, 77, 78]. Most MAGs are affiliated with filamentous taxa in the orders Synechococcales (n=17), Oscillatoriales (n=8), and Nostocales (n=4), highlighting the importance of filamentous Cyanobacteria as the ecosystem builders of polar microbial mats [4, 5, 79, 80]. Moreover, Cyanobacteria belonging to the order Nostocales often dominate the microbial communities in oligotrophic polar environments due to their ability to fix atmospheric nitrogen [4–7]. As observed previously (*e.g*. Olson *et al*. [81]), genes encoding the different subunits of the nitrogenase enzyme (NifHDK) involved in nitrogen fixation were exclusive to the four Nostocales MAGs **(Fig. 3)**.

Most Synechococcales MAGs (n=13) are phylogenetically related to strains that have been traditionally classified as *Leptolyngbya*, which is a morphological group comprising Cyanobacteria with a thin, simple filamentous morphotype that includes many different genera according to molecular data [27, 62]. Our *Leptolyngbya* MAGs can be broadly categorized into four major lineages **(Fig. 2)**: i) *Leptolyngbya stricto sensu* (‘PMM_0007’ and ‘PMM_0083’), ii) *Leptolyngbya-Stenomitos* (‘PMM_0034’ and ‘PMM_0084’), iii) *Leptolyngbya-Nodosilinea* (‘PMM_0020’ and ‘PMM_0058’), and iv) *Leptolyngbya-Oculatella-Elainella* (‘PMM_0085’, ‘PMM_0002’, ‘PMM_0041’, ‘PMM_0099’, ‘PMM_0036’, ‘PMM_0073’, and ‘PMM_0027’). The other four MAGs of filamentous Synechococcales are affiliated with the early branching *Pseudanabaena* **(Fig. 2)**. Two of these (‘PMM_0003’ and ‘PMM_0028’) are most closely related (80.9–81.0% ANI) to the strain *Pseudanabaena* sp. ULC068 isolated from a lake in the Canadian sub-Arctic (W. Vincent, unpublished) **(Table S3)**, and also clustered alongside the strains BC1403 from Greenland [28] and lw0831 from Svalbard [82] **(Fig. 2)**. The other two *Pseudanabaena* MAGs (‘PMM_0039’ and ‘PMM_0082’) are distantly related (<80% ANI) to *Synechococcus* sp. PCC 7502, a unicellular strain isolated from an alpine *Sphagnum* bog that clusters with the early branching *Pseudanabaena* [26].

Other MAGs of filamentous Cyanobacteria are affiliated with the order Oscillatoriales (n=8) **(Fig. 2)**. Four of these (‘PMM_0004’, ‘PMM_0037’, ‘PMM_0059’, and ‘PMM_0080’) are most closely related (90.1–93.2% ANI) to the strain Phorm 46 isolated from a lake in the Canadian High-Arctic [83] **(Table S3)**, and also clustered alongside strains of *Tychonema* and *Microcoleus vaginatus* **(Fig. 2)**. The other two MAGs of filamentous Oscillatoriales (‘PMM_0001’ and ‘PMM_0008’) are distantly related (<80% ANI) to the strain *Phormidium ambiguum* IAM M-71, which has an uncertain phylogenetic placement. Phylogenetic analysis of the amplified 16S rRNA gene sequence (accession AB003167) originally placed *P*. *ambiguum* IAM M-71 alongside other Oscillatoriales such as *Oscillatoria* and *Lyngbya* [84, 85]. However, a later 16S rRNA phylogeny [86] and a phylogenomic tree based on 834 single-copy genes [87] both placed the strain IAM M-71 in a similar phylogenetic position as the one inferred here, *i.e*. basal to the Nostocales **(Fig. 2)**. A BLAST analysis suggests that the AB003167 sequence is chimeric with *Phormidium muscicola* IAM M-221, but a phylogenetic artefact based on long branch attraction is also possible given the lack of related genomes. Interestingly, the *P. ambiguum* MAGs were the most widespread MAGs in our dataset, being detected in five samples in the Antarctic Peninsula, Transantarctic Mountains, and East Antarctica **(Table S4)**. Finally, the two remaining Oscillatoriales MAGs (‘PMM_0019’ and ‘PMM_0100’) clustered alongside *Crinalium epipsammum* PCC 9333 **(Fig. 2)**. *Crinalium* is a filamentous genus of Cyanobacteria with unusual elliptical trichomes [88]. Sequences related to *Crinalium* have been recovered from different alpine habitats [89] and a new species, *C*. *glaciale*, has been described from cryoconite pools in Antarctica on the basis of morphological identification [90].

In addition to filamentous taxa, we also recovered MAGs related to unicellular Cyanobacteria in the orders Gloeobacterales (n=2), Synechococcales (n=2), and Chroococcales (n=2) **(Fig. 2)**. All except the two Gloeobacterales MAGs were distantly related (<80% ANI) to genomes currently available in GenBank **(Table S3)**. The two Synechococcales MAGs (‘PMM_0022’ and ‘PMM_0072’) clustered alongside *Chamaesiphon minutus* PCC 6605 and *Chamaesiphon polymorphus* CCALA 037 **(Fig. 2)**. *Chamaesiphon* is a cosmopolitan genus that is often reported in polar and alpine terrestrial and aquatic environments, and includes two species potentially endemic to Antarctica (*C*. *arctowskii* and *C*. *austro-polonicus*) [15, 19, 20, 79, 80, 91]. Finally, the two Chroococcales MAGs (‘PMM_0021’ and ‘PMM_0086’) formed a distinct lineage related to *Microcystis* and several MAGs recovered from Arctic lakes (BioProject PRJEB38681) **(Fig. 2)**.

### Description of *Candidatus* Sivonenia alaskensis

We investigated in more detail the two Gloeobacterales MAGs ‘PMM_0042’ and ‘PMM_0068’ given the importance of this group as the most basal lineage of extant Cyanobacteria [92–94]. Phylogenetic analysis based on a concatenated alignment of 38 ribosomal proteins placed both MAGs alongside the Arctic Peat MAG ‘IMG_3300025548_6’ [39], with which they share 97.2% ANI **(Fig. 4a)**. This is above the threshold of 95–96% ANI commonly used for delineating microbial species [95, 96], which thus suggests that the three MAGs (‘PMM_0042’, ‘PMM_0068’, and ‘IMG_3300025548_6’) belong to the same species. Furthermore, their phylogenetic placement and low ANI (<80%) with other Gloeobacterales indicate that they constitute a distinct genus in this order. Separation from the other Gloeobacterales is also supported by analysis of the 16S rRNA gene of the MAG ‘IMG_3300025548_6’, which is 91.8–92.0%, 90.0%, and 89.3% similar to the sequences of *Gloeobacter* spp., *Candidatus* Aurora vandensis, and *Anthocerotibacter panamensis*, respectively **(Fig. 4b)**. We consider that the MAGs ‘PMM_0042’, ‘PMM_0068’, and ‘IMG_3300025548_6’ represent a novel lineage in the order Gloeobacterales and propose the name *Candidatus* Sivonenia alaskensis (*Sivonenia*: in honour of our colleague and Cyanobacteria expert Dr. Kaarina Sivonen, *professor emerita* of the University of Helsinki; *alaskensis*: relative to the geographic origin of the MAG ‘IMG_3300025548_6’, which is proposed here as the nomenclatural type for this species according to the SeqCode initiative [97]).

**Fig. 4.**
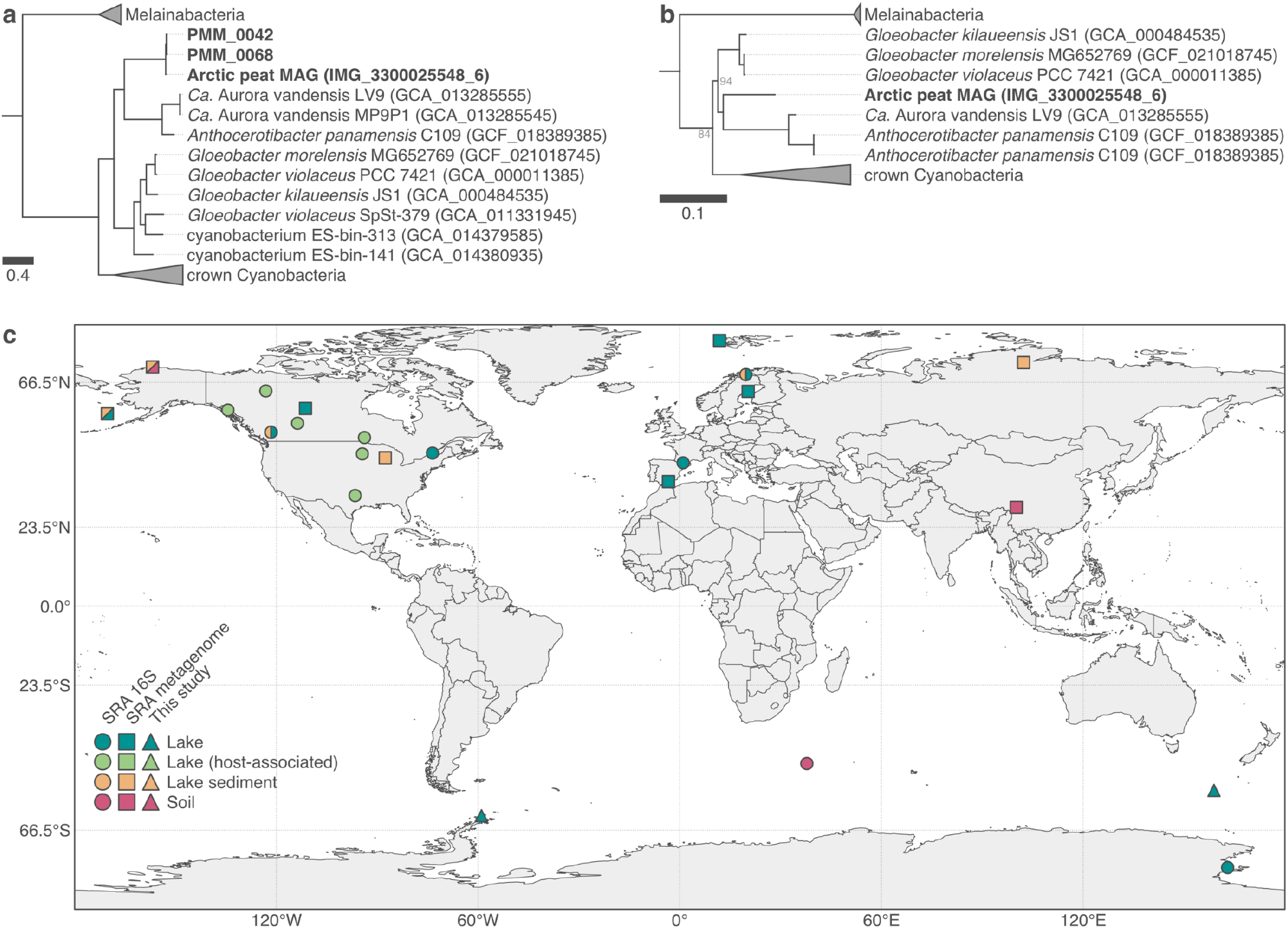
*Candidatus* Sivonenia alaskensis, a lineage of early branching Cyanobacteria in the order Gloeobacterales. **a)** Maximum-likelihood tree (LG+R8 model) based on a concatenated alignment of 38 ribosomal proteins from the three *Ca*. Sivonenia alaskensis MAGs (in bold) and other Gloeobacterales and selected genomes from GenBank. All nodes have bootstrap support ≥95%. **b)** Maximum-likelihood tree (GTR+F+R8 model) of the 16S rRNA gene of *Ca*. Sivonenia alaskensis. Nodes have bootstrap support ≥95% unless shown otherwise. **c)** Geographic distribution of *Ca*. Sivonenia alaskensis based on significant matches with metagenomic and 16S rRNA gene amplicon sequencing datasets in SRA (≥20% containment and ≥0.1% relative abundance, respectively).

### *In silico* analysis indicates that *Ca*. Sivonenia alaskensis is a thylakoid-less cyanobacterium

Analysis of the protein-coding genes of the *Ca*. Sivonenia alaskensis MAGs revealed many similarities with other Gloeobacterales, thus supporting their phylogenetic placement within this order of early branching Cyanobacteria **(Fig. 4ab)**. For instance, strains of *Gloeobacter* spp. and *A*. *panamensis* differ notoriously from other Cyanobacteria by the lack of thylakoid membranes and the presence of a reduced photosynthetic apparatus [98–101]. These traits are considered ancestral features of oxygenic photosynthesis given the basal position of Gloeobacterales in the evolution of Cyanobacteria and plastids [92–94]. Like the genomes of other Gloeobacterales [39, 101–104], the *Ca*. Sivonenia alaskensis MAGs lack the genes for several subunits of the photosystems I (PsaI, PsaJ, PsaK, and PsaX) and II (PsbY, PsbZ, and Psb27), the circadian clock (KaiA, KaiB, and KaiC), and the thylakoid membrane (SqdB and SqdX) **(Fig. 3, Table S5)**. Moreover, similarly to *A*. *panamensis* and *Ca*. Aurora vandensis but unlike *Gloeobacter* spp. [101], *Ca*. Sivonenia alaskensis lacks two subunits of the photosystem II (PsbM and PsbU) and the cytochrome *b_6_f* complex (PetM and PetN), and does not contain any gene involved in the synthesis of phycoerythrin (Pec). Overall, the *in silico* analysis of the proteome of *Ca*. Sivonenia alaskensis suggests that this lineage comprises organisms without thylakoid membranes and with a reduced photosynthetic machinery, both of which are the defining characteristics of the order Gloeobacterales [62]. Moreover, the predicted structure of the photosystem II, cytochrome *b_6_f*, and phycobilisome machineries of *Ca*. Sivonenia alaskensis holds more similarities with *A*. *panamensis* and *Ca*. Aurora vandensis than with *Gloeobacter* spp., supporting the evolutionary relationship inferred from the analysis of ribosomal proteins and the 16S rRNA gene **(Fig. 4ab)**. Whether the unique characteristics of *Gloeobacter* spp. reflect the ancestral state of the phylum Cyanobacteria has been an open question in the study of the early evolution of this group for many decades. The discovery of *Ca*. Sivonenia alaskensis, *A*. *panamensis* [101], and *Ca*. Aurora vandensis [40] suggests that characteristics such as the lack of thylakoid membranes and a reduced photosynthetic apparatus are indeed a defining characteristic of the early branching Gloeobacterales.

### *Ca*. Sivonenia alaskensis is predominantly distributed in the cold biosphere

Read recruitment analysis revealed that the two *Ca*. Sivonenia alaskensis MAGs are found in four microbial mat samples from the sub-Antarctic and Antarctic Peninsula, where they constitute up to 1.0% of the metagenomes **(Table S4)**. To gain further insights into the ecology of *Ca*. Sivonenia alaskensis, we used *sourmash branchwater* [69, 70] to search metagenomic datasets in SRA for sequences matching the *Ca*. Sivonenia alaskensis MAGs. We also searched its 16S rRNA gene in amplicon sequencing datasets in SRA using IMNGS [71]. This extensive search, which included collectively *ca*. 1.3 million public datasets from around the globe, revealed sequences related to *Ca*. Sivonenia alaskensis in lakes, sediments, and soils mainly in polar, sub-polar, and alpine environments **(Fig. 4c)**. Sequences matching the *Ca*. Sivonenia alaskensis MAGs were particularly abundant (0.4–8.0%) in amplicon sequencing datasets of active communities (*i.e*. derived from RNA molecules) in the sediment of thermokarst lakes near Barrow (Alaska) [105], as well as in metagenomic datasets from the sediment of Lake Hill (St. Paul Island, Alaska) [106] (0.8–3.1% of the reads). Interestingly, sequences matching the 16S rRNA gene of *Ca*. Sivonenia alaskensis were found in several datasets obtained from the gut microbiome of stickleback fishes (Actinopterygii: Gasterosteidae) and mayflies (Insecta: Ephemeroptera) [107–109].

Despite their importance for the study of the evolution of oxygenic photosynthesis, little is known about the ecology of the early branching Gloeobacterales compared to the other Cyanobacteria [83, 92, 101, 110]. *Gloeobacter*, which was for many decades the only described genus in this order, is typically found in low light, wet rock habitats [98, 111, 112]. Amplicon sequencing studies have also reported 16S rRNA gene sequences loosely related to *Gloeobacter* spp. in Arctic [17, 21] and temperate [113] soil crusts, and in Arctic [78, 114] and Antarctic [115] microbial mats. The phylogenetic and ecological range of the order Gloeobacterales has expanded recently with the discovery of *Ca*. Aurora vandensis from Antarctic lakes [40], *A*. *panamensis* associated with a tropical bryophyte [101], and five MAGs from different ecosystems including the *Ca*. Sivonenia alaskensis MAG ‘IMG_3300025548_6’ recovered from Arctic peat soil [39]. Apart from *A*. *panamensis*, Gloeobacterales appear to show a preference for low light and cold environments. This has been linked to their slow growth which, in turn, appears to be a consequence of their reduced photosynthetic apparatus [101]. In agreement with this, our results suggest that *Ca*. Sivonenia alaskensis is predominant in cold regions, especially polar and alpine lakes and sediments **(Fig. 4c)**. Its high abundance in an RNA-derived amplicon sequencing dataset of lake sediments in Alaska [105] suggests that *Ca*. Sivonenia alaskensis forms active populations in this habitat. By contrast, the detection of sequences matching the 16S rRNA gene of *Ca*. Sivonenia alaskensis in the microbiome of stickleback fishes does not entail that they are active members of gut communities. These sequences likely represent cells that were ingested either incidentally or collaterally via zooplankton that is consumed by the fish (Daniel Bolnick, personal communication).

### Analysis of resistance mechanisms to environmental stress in *Ca*. Sivonenia alaskensis

To obtain insights regarding the distribution of *Ca*. Sivonenia alaskensis across the cold biosphere, we searched the MAGs for genes involved in resistance mechanisms to environmental stress. We found 75 genes related to mechanisms to cope with desiccation, cold, and ultraviolet radiation (UVR) stresses in at least one of the *Ca*. Sivonenia alaskensis MAGs **(Table S6)**. Among these are genes involved in the Wzy- and ABC transporter-dependent pathways for the assembly and export of extracellular polymeric substances (EPS). The production of an EPS matrix is a mechanism that is commonly employed by Cyanobacteria to cope with desiccation and freezing [116]. Genes involved in the synthase-dependent pathway of EPS production were not found. Scytonemin and mycosporine-like amino acids (MAAs) are often produced by Cyanobacteria as UVR-screening compounds [117]. Despite having several of the genes involved in the production of scytonemin and MAAs, the genes encoding the key proteins ScyC, ScyD, EboA, EboB, EboC, and MysC were not found. As such, the production of these compounds by *Ca*. Sivonenia alaskensis is unlikely. We identified several mechanisms of resistance to cold in the *Ca*. Sivonenia alaskensis MAGs, including proteins involved in the regulation of the cell membrane fluidity, regulation of replication and translation, and RNA metabolism **(Table S6)**. Finally, mechanisms of DNA repair include the base excision repair pathway for several glycosylases, the homologous recombination pathway for single-stranded breaks, and one of the subtypes of the nuclear excision repair pathway.

## Conclusion

We investigated 17 polar microbial mat metagenomes and recovered 37 MAGs of Cyanobacteria representing different levels of phylogenetic novelty: around half of the MAGs are very distant (<80% ANI) to genomes currently available in GenBank; the other half are related to polar and alpine strains with varying levels of genome similarity (80.1–99.8% ANI). Among the latter, we describe the phylogenetic, metabolic potential, and ecological characteristics of a lineage in the early branching Gloeobacterales. *In silico* analyses indicate that this lineage – which we name *Ca*. Sivonenia alaskensis – is a thylakoid-less cyanobacterium that is mostly found in cold environments and harbours common mechanisms of resistance to environmental stress. Our study shows that genome-resolved metagenomics is a reliable and straightforward way of recovering novel genomes of Cyanobacteria without the need for strain isolation. However, strain isolation is still useful for many purposes and may in fact benefit from genomic information obtained from MAGs to design protocols for targeted isolation. Based on the ANI thresholds commonly used for delineating microbial species [95, 96], most of the MAGs obtained represent different species or even genera from the ones currently represented by genomes in GenBank. Comparison with strains without genome data was not possible as only one of the 37 MAGs included the 16S rRNA gene, which is the most widely used molecular marker for the taxonomy of Cyanobacteria [27]. The use of long read technologies (*e.g*. Oxford Nanopore and PacBio SMRT sequencing) could help alleviate this issue. Altogether, our study highlights the uniqueness of polar microbiomes and their specialized communities of Cyanobacteria, of which a large fraction is yet to be characterized.

## Author statements

### Author contributions

ISP, YL, EV, and AW conceived the experiments. ISP performed most of the analysis, and RVP, VS, and BRR contributed with minor parts. ISP and RVP wrote the manuscript. All authors provided important feedback, helped shape the study, and contributed to the writing of the manuscript.

## Acknowledgments

The authors would like to acknowledge Sofie D’Hondt (UGent) and Bjorn Tytgat for help with DNA extraction and library preparation, the IT Centre for Science – CSC (Finland) for providing the computational resources used in the study, and Kaarina Sivonen, Daniel Bolnick, Danillo Alvarenga, and Tânia Shishido for comments. We also thank Dominic A. Hodgson, Steve J. Roberts, Wim Van Nieuwenhuyze, Koen Sabbe, Dagmar Obbels, Otakar Strunecký, Kate Kopalová, Jan Kavan, Josef Elster, Pieter Vanormelingen, and Eveline Pinseel for help during sampling campaigns or/and sharing samples.

## References

1. Golubic S. Modern stromatolites: a review. In: Riding R (editor). Calcareous Algae and Stromatolites. Berlin, Heidelberg: Springer Berlin Heidelberg. pp. 541–561.

2. Stal LJ. Cyanobacterial mats and stromatolites. In: Whitton BA (editor). Ecology of Cyanobacteria II. Dordrecht: Springer Netherlands. pp. 65–125.

3. Bolhuis H, Cretoiu MS, Stal LJ. Molecular ecology of microbial mats. FEMS Microbiol Ecol 2014;90:335–350.

4. Singh SM, Elster J. Cyanobacteria in Antarctic lake environments: a mini-review. In: Seckbach J (editor). Algae and Cyanobacteria in Extreme Environments. Dordrecht: Springer Netherlands. pp. 303–320.

5. Vincent WF, Quesada A. Cyanobacteria in high latitude lakes, rivers and seas. In: Whitton BA (editor). Ecology of Cyanobacteria II. Dordrecht: Springer Netherlands. pp. 371–385.

6. Quesada A, Vincent WF. Cyanobacteria in the cryosphere: snow, ice and extreme cold. In: Whitton BA (editor). Ecology of Cyanobacteria II. Dordrecht: Springer Netherlands. pp. 387–399.

7. Van Goethem MW, Cowan DA. Role of Cyanobacteria in the ecology of polar environments. In: Castro-Sowinski S (editor). The Ecological Role of Micro-organisms in the Antarctic Environment. Cham: Springer International Publishing. pp. 3–23.

8. Chrismas NAM, Anesio AM, Sánchez-Baracaldo P. The future of genomics in polar and alpine cyanobacteria. FEMS Microbiol Ecol 2018;94:fiy032.

9. Vincent WF, Downes MT, Castenholz RW, Howard-Williams C. Community structure and pigment organisation of cyanobacteria-dominated microbial mats in Antarctica. Eur J Phycol 1993;28:213–221.

10. Elster J, Komarek O. Ecology of periphyton in a meltwater stream ecosystem in the maritime Antarctic. Antarct Sci 2003;15:189–201.

11. Elster J, Lukesová A, Svoboda J, Kopecky J, Kanda H. Diversity and abundance of soil algae in the polar desert, Sverdrup Pass, central Ellesmere Island. Polar Rec 1999;35:231–254.

12. Taton A, Grubisic S, Ertz D, Hodgson DA, Piccardi R, et al. Polyphasic study of Antarctic cyanobacterial strains. J Phycol 2006;42:1257–1270.

13. Palinska KA, Schneider T, Surosz W. Phenotypic and phylogenetic studies of benthic mat-forming cyanobacteria on the NW Svalbard. Polar Biol 2017;40:1607–1616.

14. Strunecky O, Raabova L, Bernardova A, Ivanova AP, Semanova A, et al. Diversity of cyanobacteria at the Alaska North Slope with description of two new genera: *Gibliniella* and *Shackletoniella*. FEMS Microbiol Ecol 2020;96:fiz189.

15. Taton A, Hoffmann L, Wilmotte A. Cyanobacteria in microbial mats of Antarctic lakes (East Antarctica): a microscopical approach. Algol Stud 2008;126:173–208.

16. Namsaraev Z, Mano M-J, Fernandez R, Wilmotte A. Biogeography of terrestrial cyanobacteria from Antarctic ice-free areas. Ann Glaciol 2010;51:171–177.

17. Pushkareva E, Pessi IS, Wilmotte A, Elster J. Cyanobacterial community composition in Arctic soil crusts at different stages of development. FEMS Microbiol Ecol 2015;91:fiv143.

18. Pushkareva E, Pessi IS, Namsaraev Z, Mano M-J, Elster J, et al. Cyanobacteria inhabiting biological soil crusts of a polar desert: Sør Rondane Mountains, Antarctica. Syst Appl Microbiol 2018;41:363–373.

19. Pessi IS, Maalouf PDC, Laughinghouse HD, Baurain D, Wilmotte A. On the use of high-throughput sequencing for the study of cyanobacterial diversity in Antarctic aquatic mats. J Phycol 2016;52:356–368.

20. Pessi IS, Lara Y, Durieu B, Maalouf P de C, Verleyen E, et al. Community structure and distribution of benthic cyanobacteria in Antarctic lacustrine microbial mats. FEMS Microbiol Ecol 2018;94:fiy042.

21. Pessi IS, Pushkareva E, Lara Y, Borderie F, Wilmotte A, et al. Marked succession of cyanobacterial communities following glacier retreat in the High Arctic. Microb Ecol 2019;77:136–147.

22. Taton A, Grubisic S, Brambilla E, De Wit R, Wilmotte A. Cyanobacterial diversity in natural and artificial microbial mats of Lake Fryxell (McMurdo Dry Valleys, Antarctica): a morphological and molecular approach. Appl Environ Microbiol 2003;69:5157–5169.

23. Taton A, Grubisic S, Balthasart P, Hodgson DA, Laybourn-Parry J, et al. Biogeographical distribution and ecological ranges of benthic cyanobacteria in East Antarctic lakes. FEMS Microbiol Ecol 2006;57:272–289.

24. Fernandez-Carazo R, Hodgson DA, Convey P, Wilmotte A. Low cyanobacterial diversity in biotopes of the Transantarctic Mountains and Shackleton Range (80–82°S), Antarctica. FEMS Microbiol Ecol 2011;77:503–517.

25. Wilmotte A, Golubić S. Morphological and genetic criteria in the taxonomy of Cyanophyta/Cyanobacteria. Algol Stud 1991;64:1–24.

26. Shih PM, Wu D, Latifi A, Axen SD, Fewer DP, et al. Improving the coverage of the cyanobacterial phylum using diversity-driven genome sequencing. Proc Natl Acad Sci 2013;110:1053–1058.

27. Mareš J. Multilocus and SSU rRNA gene phylogenetic analyses of available cyanobacterial genomes, and their relation to the current taxonomic system. Hydrobiologia 2018;811:19–34.

28. Chrismas NAM, Barker G, Anesio AM, Sánchez-Baracaldo P. Genomic mechanisms for cold tolerance and production of exopolysaccharides in the Arctic cyanobacterium *Phormidesmis priestleyi* BC1401. BMC Genomics 2016;17:533.

29. Péquin B, Tremblay J, Maynard C, Wasserscheid J, Greer CW. Draft whole-genome sequences of the polar cyanobacterium *Leptolyngbya* sp. strain Cla-17 and its associated flavobacterium. Microbiol Resour Announc 2022;11:e00059–22.

30. Lara Y, Durieu B, Cornet L, Verlaine O, Rippka R, et al. Draft genome sequence of the axenic strain *Phormidesmis priestleyi* ULC007, a cyanobacterium isolated from Lake Bruehwiler (Larsemann Hills, Antarctica). Genome Announc 2017;5:e01546–16.

31. Chrismas NAM, Williamson CJ, Yallop ML, Anesio AM, Sánchez-Baracaldo P. Photoecology of the Antarctic cyanobacterium *Leptolyngbya* sp. BC1307 brought to light through community analysis, comparative genomics and in vitro photophysiology. Mol Ecol 2018;27:5279–5293.

32. Tang J, Du L-M, Liang Y-M, Daroch M. Complete genome sequence and comparative analysis of *Synechococcus* sp. CS-601 (SynAce01), a cold-adapted cyanobacterium from an oligotrophic Antarctic habitat. Int J Mol Sci 2019;20:152.

33. Effendi DB, Sakamoto T, Ohtani S, Awai K, Kanesaki Y. Possible involvement of extracellular polymeric substrates of Antarctic cyanobacterium *Nostoc* sp. strain SO-36 in adaptation to harsh environments. J Plant Res 2022;135:771–784.

34. Cornet L, Bertrand AR, Hanikenne M, Javaux EJ, Wilmotte A, et al. Metagenomic assembly of new (sub)polar Cyanobacteria and their associated microbiome from non-axenic cultures. Microb Genomics 2018;4:212.

35. Chen L-X, Anantharaman K, Shaiber A, Eren AM, Banfield JF. Accurate and complete genomes from metagenomes. Genome Res 2020;30:315–333.

36. Delmont TO. Discovery of nondiazotrophic *Trichodesmium* species abundant and widespread in the open ocean. Proc Natl Acad Sci 2021;118:e2112355118.

37. Pessi IS, Viitamäki S, Virkkala A-M, Eronen-Rasimus E, Delmont TO, et al. In-depth characterization of denitrifier communities across different soil ecosystems in the tundra. Environ Microbiome 2022;17:30.

38. Pessi IS, Rutanen A, Hultman J. *Candidatus* Nitrosopolaris, a genus of putative ammonia-oxidizing archaea with a polar/alpine distribution. FEMS Microbes 2022;3:xtac019.

39. Grettenberger CL. Novel *Gloeobacterales* spp. from diverse environments across the globe. mSphere 2021;6:e00061–21.

40. Grettenberger CL, Sumner DY, Wall K, Brown CT, Eisen JA, et al. A phylogenetically novel cyanobacterium most closely related to *Gloeobacter*. ISME J 2020;14:2142–2152.

41. Lumian JE, Jungblut AD, Dillion ML, Hawes I, Doran PT, et al. Metabolic capacity of the Antarctic cyanobacterium *Phormidium pseudopriestleyi* that sustains oxygenic photosynthesis in the presence of hydrogen sulfide. Genes 2021;12:426.

42. Terauds A, Lee JR. Antarctic biogeography revisited: updating the Antarctic Conservation Biogeographic Regions. Divers Distrib 2016;22:836–840.

43. Ewels P, Magnusson M, Lundin S, Käller M. MultiQC: summarize analysis results for multiple tools and samples in a single report. Bioinformatics 2016;32:3047–3048.

44. Martin M. Cutadapt removes adapter sequences from high-throughput sequencing reads. EMBnet.journal 2011;17:10.

45. Bengtsson-Palme J, Hartmann M, Eriksson KM, Pal C, Thorell K, et al. METAXA2: improved identification and taxonomic classification of small and large subunit rRNA in metagenomic data. Mol Ecol Resour 2015;15:1403–1414.

46. Schloss PD, Westcott SL, Ryabin T, Hall JR, Hartmann M, et al. Introducing mothur: open-source, platform-independent, community-supported software for describing and comparing microbial communities. Appl Environ Microbiol 2009;75:7537–7541.

47. Quast C, Pruesse E, Yilmaz P, Gerken J, Schweer T, et al. The SILVA ribosomal RNA gene database project: improved data processing and web-based tools. Nucleic Acids Res 2013;41:D590–D596.

48. Wang Q, Garrity GM, Tiedje JM, Cole JR. Naïve Bayesian classifier for rapid assignment of rRNA sequences into the new bacterial taxonomy. Appl Environ Microbiol 2007;73:5261–5267.

49. Li D, Liu C-M, Luo R, Sadakane K, Lam T-W. MEGAHIT: an ultra-fast single-node solution for large and complex metagenomics assembly via succinct de Bruijn graph. Bioinformatics 2015;31:1674–1676.

50. Eren AM, Kiefl E, Shaiber A, Veseli I, Miller SE, et al. Community-led, integrated, reproducible multi-omics with anvi’o. Nat Microbiol 2021;6:3–6.

51. Hyatt D, Chen G-L, LoCascio PF, Land ML, Larimer FW, et al. Prodigal: prokaryotic gene recognition and translation initiation site identification. BMC Bioinformatics 2010;11:119.

52. Eddy SR. Accelerated profile HMM searches. PLoS Comput Biol 2011;7:e1002195.

53. Lee MD. GToTree: a user-friendly workflow for phylogenomics. Bioinformatics 2019;35:4162–4164.

54. Buchfink B, Xie C, Huson DH. Fast and sensitive protein alignment using DIAMOND. Nat Methods 2015;12:59–60.

55. Parks DH, Chuvochina M, Rinke C, Mussig AJ, Chaumeil P-A, et al. GTDB: an ongoing census of bacterial and archaeal diversity through a phylogenetically consistent, rank normalized and complete genome-based taxonomy. Nucleic Acids Res 2022;50:D785–D794.

56. Langmead B, Salzberg SL. Fast gapped-read alignment with Bowtie 2. Nat Methods 2012;9:357–359.

57. Li H, Handsaker B, Wysoker A, Fennell T, Ruan J, et al. The Sequence Alignment/Map format and SAMtools. Bioinformatics 2009;25:2078–2079.

58. Chaumeil P-A, Mussig AJ, Hugenholtz P, Parks DH. GTDB-Tk: a toolkit to classify genomes with the Genome Taxonomy Database. Bioinformatics 2020;36:1925–1927.

59. Edgar RC. MUSCLE: multiple sequence alignment with high accuracy and high throughput. Nucleic Acids Res 2004;32:1792–1797.

60. Nguyen L-T, Schmidt HA, von Haeseler A, Minh BQ. IQ-TREE: a fast and effective stochastic algorithm for estimating maximum-likelihood phylogenies. Mol Biol Evol 2015;32:268–274.

61. Jain C, Rodriguez-R LM, Phillippy AM, Konstantinidis KT, Aluru S. High throughput ANI analysis of 90K prokaryotic genomes reveals clear species boundaries. Nat Commun 2018;9:5114.

62. Komárek J, Kaštovský J, Mareš J, Johansen JR. Taxonomic classification of cyanoprokaryotes (cyanobacterial genera) 2014, using a polyphasic approach. Preslia 2014;86:295–335.

63. Aramaki T, Blanc-Mathieu R, Endo H, Ohkubo K, Kanehisa M, et al. KofamKOALA: KEGG ortholog assignment based on profile HMM and adaptive score threshold. Bioinformatics 2020;36:2251–2252.

64. Mistry J, Chuguransky S, Williams L, Qureshi M, Salazar GA, et al. Pfam: the protein families database in 2021. Nucleic Acids Res 2021;49:D412–D419.

65. Galperin MY, Wolf YI, Makarova KS, Vera Alvarez R, Landsman D, et al. COG database update: focus on microbial diversity, model organisms, and widespread pathogens. Nucleic Acids Res 2021;49:D274–D281.

66. Pearson WR. An introduction to sequence similarity (“homology”) searching. Curr Protoc Bioinforma 2013;42:3.1.1–3.1.8.

67. Olm MR, Brown CT, Brooks B, Banfield JF. dRep: a tool for fast and accurate genomic comparisons that enables improved genome recovery from metagenomes through dereplication. ISME J 2017;11:2864–2868.

68. Li H. Minimap and miniasm: fast mapping and de novo assembly for noisy long sequences. Bioinformatics 2016;32:2103–2110.

69. Irber L, Pierce-Ward NT, Brown CT. Sourmash branchwater enables lightweight petabyte-scale sequence search. Preprint (BioRxiv). Epub ahead of print 3 November 2022. DOI: 10.1101/2022.11.02.514947.

70. Lumian J, Sumner D, Grettenberger C, Jungblut AD, Irber L, et al. Biogeographic distribution of five Antarctic cyanobacteria using large-scale k-mer searching with sourmash branchwater. Preprint (BioRxiv). Epub ahead of print 30 October 2022. DOI: 10.1101/2022.10.27.514113.

71. Lagkouvardos I, Joseph D, Kapfhammer M, Giritli S, Horn M, et al. IMNGS: a comprehensive open resource of processed 16S rRNA microbial profiles for ecology and diversity studies. Sci Rep 2016;6:33721.

72. Zaikova E, Goerlitz DS, Tighe SW, Wagner NY, Bai Y, et al. Antarctic relic microbial mat community revealed by metagenomics and metatranscriptomics. Front Ecol Evol 2019;7:1.

73. Slattery M, Lesser MP. Allelopathy-mediated competition in microbial mats from Antarctic lakes. FEMS Microbiol Ecol 2017;93:fix019.

74. Varin T, Lovejoy C, Jungblut AD, Vincent WF, Corbeil J. Metagenomic analysis of stress genes in microbial mat communities from Antarctica and the High Arctic. Appl Environ Microbiol 2012;78:549–559.

75. Soo RM, Skennerton CT, Sekiguchi Y, Imelfort M, Paech SJ, et al. An expanded genomic representation of the phylum Cyanobacteria. Genome Biol Evol 2014;6:1031–1045.

76. Komárek J, Genuário DB, Fiore MF, Elster J. Heterocytous cyanobacteria of the Ulu Peninsula, James Ross Island, Antarctica. Polar Biol 2015;38:475–492.

77. Chrismas NAM, Anesio AM, Sánchez-Baracaldo P. Multiple adaptations to polar and alpine environments within cyanobacteria: a phylogenomic and Bayesian approach. Front Microbiol moo;6:1070.

78. Jungblut AD, Lovejoy C, Vincent WF. Global distribution of cyanobacterial ecotypes in the cold biosphere. ISME J 2010;4:191–202.

79. Velichko N, Smirnova S, Averina S, Pinevich A. A survey of Antarctic cyanobacteria. Hydrobiologia 2021;848:2627–2652.

80. Davydov D. Cyanobacterial diversity of Svalbard Archipelago. Polar Biol 2021;44:1967–1978.

81. Olson JB, Steppe TF, Litaker RW, Paerl HW. N2-fixing microbial consortia associated with the ice cover of Lake Bonney, Antarctica. Microb Ecol 1998;36:231–238.

82. Su H-N, Wang Q-M, Li C-Y, Li K, Luo W, et al. Structural insights into the cold adaptation of the photosynthetic pigment-protein C-phycocyanin from an Arctic cyanobacterium. Biochim Biophys Acta BBA - Bioenerg 2017;1858:325–335.

83. Moore KR, Magnabosco C, Momper L, Gold DA, Bosak T, et al. An expanded ribosomal phylogeny of Cyanobacteria supports a deep placement of plastids. Front Microbiol 2019;10:1612.

84. Ishida T, Yokota A, Sugiyama J. Phylogenetic relationships of filamentous cyanobacterial taxa inferred from 16S rRNA sequence divergence. J Gen Appl Microbiol 1997;43:237–241.

85. Honda D, Yokota A, Sugiyama J. Detection of seven major evolutionary lineages in cyanobacteria based on the 16S rRNA gene sequence analysis with new sequences of five marine *Synechococcus* strains. J Mol Evol 1999;48:723–739.

86. Ishida T, Watanabe MM, Sugiyama J, Yokota A. Evidence for polyphyletic origin of the members of the orders of Oscillatoriales and Pleurocapsales as determined by 16S rDNA analysis. FEMS Microbiol Lett 2001;201:79–82.

87. Chen M-Y, Teng W-K, Zhao L, Hu C-X, Zhou Y-K, et al. Comparative genomics reveals insights into cyanobacterial evolution and habitat adaptation. ISME J 2021;15:211–227.

88. de Winder B, Stal LJ, Mur LR. *Crinalium epipsammum* sp. nov.: a filamentous cyanobacterium with trichomes composed of elliptical cells and containing poly-ß-(1,4) glucar (cellulose). J Gen Microbiol 1990;136:1645–1653.

89. Bohunická M, Mareš J, Hrouzek P, Urajová P, Lukeš M, et al. A combined morphological, ultrastructural, molecular, and biochemical study of the peculiar family Gomontiellaceae (Oscillatoriales) reveals a new cylindrospermopsin-producing clade of cyanobacteria. J Phycol 2015;51:1040–1054.

90. Broady PA, Kibblewhite AL. Morphological characterization of Oscillatoriales (Cyanobacteria) from Ross Island and southern Victoria Land, Antarctica. Antarct Sci 1991;3:35–45.

91. Komárek J. Phenotypic and ecological diversity of freshwater coccoid cyanobacteria from maritime Antarctica and Islands of NW Weddell Sea. II. Czech Polar Rep 2014;4:17–39.

92. Demoulin CF, Lara YJ, Cornet L, François C, Baurain D, et al. Cyanobacteria evolution: insight from the fossil record. Free Radic Biol Med 2019;140:206–223.

93. Seo P-S, Yokota A. The phylogenetic relationships of cyanobacteria inferred from 16S rRNA, *gyrB, rpoC1* and *rpoD1* gene sequences. J Gen Appl Microbiol 2003;49:191–203.

94. Gupta RS, Mathews DW. Signature proteins for the major clades of Cyanobacteria. BMC Evol Biol 2010;10:24.

95. Konstantinidis KT, Rosselló-Móra R, Amann R. Uncultivated microbes in need of their own taxonomy. ISME J 2017;11:2399–2406.

96. Ciufo S, Kannan S, Sharma S, Badretdin A, Clark K, et al. Using average nucleotide identity to improve taxonomic assignments in prokaryotic genomes at the NCBI. Int J Syst Evol Microbiol 2018;68:2386–2392.

97. Hedlund BP, Chuvochina M, Hugenholtz P, Konstantinidis KT, Murray AE, et al. SeqCode: a nomenclatural code for prokaryotes described from sequence data. Nat Microbiol 2022;7:1702–1708.

98. Rippka R, Waterbury J, Cohen-Bazire G. A cyanobacterium which lacks thylakoids. Arch Microbiol 1974;100:419–436.

99. Guglielmi G, Cohen-Bazire G, Bryant DA. The structure of *Gloeobacter violaceus* and its phycobilisomes. Arch Microbiol 1981;129:181–189.

100. Bryant DA, Cohen-Bazire G, Glazer AN. Characterization of the biliproteins of *Gloeobacter violaceus*. Arch Microbiol 1981;129:190–198.

101. Rahmatpour N, Hauser DA, Nelson JM, Chen PY, Villarreal A. JC, et al. A novel thylakoid-less isolate fills a billion-year gap in the evolution of Cyanobacteria. Curr Biol 2021;31:2857–2867.

102. Nakamura Y. Complete genome structure of *Gloeobacter violaceus* PCC 7421, a cyanobacterium that lacks thylakoids. DNA Res 2003;10:137–145.

103. Saw JH, Cardona T, Montejano G. Complete genome sequencing of a novel *Gloeobacter* species from a waterfall cave in Mexico. Genome Biol Evol 2021;13:evab264.

104. Saw JHW, Schatz M, Brown MV, Kunkel DD, Foster JS, et al. Cultivation and complete genome sequencing of *Gloeobacter kilaueensis* sp. nov., from a lava cave in Kīlauea Caldera, Hawai’i. PLoS ONE 2013;8:e76376.

105. Matheus Carnevali PB, Herbold CW, Hand KP, Priscu JC, Murray AE. Distinct microbial assemblage structure and archaeal diversity in sediments of Arctic thermokarst lakes differing in methane sources. Front Microbiol 2018;9:1192.

106. Graham RW, Belmecheri S, Choy K, Culleton BJ, Davies LJ, et al. Timing and causes of mid-Holocene mammoth extinction on St. Paul Island, Alaska. Proc Natl Acad Sci 2016;113:9310–9314.

107. Bolnick DI, Snowberg LK, Caporaso JG, Lauber C, Knight R, et al. Major Histocompatibility Complex class IIb polymorphism influences gut microbiota composition and diversity. Mol Ecol 2014;23:4831–4845.

108. Bolnick DI, Snowberg LK, Hirsch PE, Lauber CL, Knight R, et al. Individuals’ diet diversity influences gut microbial diversity in two freshwater fish (threespine stickleback and Eurasian perch). Ecol Lett 2014;17:979–987.

109. Bolnick DI, Snowberg LK, Hirsch PE, Lauber CL, Org E, et al. Individual diet has sex-dependent effects on vertebrate gut microbiota. Nat Commun 2014;5:4500.

110. Delwiche CF. Microbial biodiversity: a newly isolated cyanobacterium sheds light on the evolution of photosynthesis. Curr Biol 2021;31:R843–R845.

111. Mareš J, Hrouzek P, Kaňa R, Ventura S, Strunecký O, et al. The primitive thylakoid-less cyanobacterium *Gloeobacter* is a common rock-dwelling organism. PLoS ONE 2013;8:e66323.

112. Golubic S, Campbell SE. Analogous microbial forms in recent subaerial habitats and in Precambrian cherts: *Gloethece coerulea* Geitler and *Eosynechococcus moorei Hofmann*. Precambrian Res 1979;8:201–217.

113. Williams L, Loewen-Schneider K, Maier S, Büdel B. Cyanobacterial diversity of western European biological soil crusts along a latitudinal gradient. FEMS Microbiol Ecol 2016;92:fiw157.

114. Lionard M, Péquin B, Lovejoy C, Vincent WF. Benthic cyanobacterial mats in the High Arctic: multi-layer structure and fluorescence responses to osmotic stress. Front Microbiol 2012;3:140.

115. Nakai R, Abe T, Baba T, Imura S, Kagoshima H, et al. Microflorae of aquatic moss pillars in a freshwater lake, East Antarctica, based on fatty acid and 16S rRNA gene analyses. Polar Biol 2012;35:425–433.

116. Pereira SB, Mota R, Vieira CP, Vieira J, Tamagnini P. Phylum-wide analysis of genes/proteins related to the last steps of assembly and export of extracellular polymeric substances (EPS) in cyanobacteria. Sci Rep 2015;5:14835.

117. Gao Q, Garcia-Pichel F. Microbial ultraviolet sunscreens. Nat Rev Microbiol 2011;9:791–802.

